# GraphCompass: Spatial metrics for differential analyses of cell organization across conditions

**DOI:** 10.1101/2024.02.02.578605

**Authors:** Mayar Ali, Merel Kuijs, Soroor Hediyeh-zadeh, Tim Treis, Karin Hrovatin, Giovanni Palla, Anna C. Schaar, Fabian J. Theis

**Affiliations:** Institute of Computational Biology, Helmholtz Munich, Germany; Institute for Tissue Engineering and Regenerative Medicine, Helmholtz Munich, Germany; Department of Mathematics, School of Computation, Information and Technology, Technical University of Munich, Germany; Graduate School of Systemic Neurosciences, Ludwig Maximilian University of Munich, Germany; School of Life Sciences, Technical University of Munich, Germany; Munich Center for Machine Learning, Technical University of Munich, Germany

**Keywords:** Spatial omics, Tissue architecture, Cell spatial organization, Graph analytics

## Abstract

Spatial omics technologies are increasingly leveraged to characterize how disease disrupts tissue organization and cellular niches. While multiple methods to analyze spatial variation within a sample have been published, statistical and computational approaches to compare cell spatial organization across samples or conditions are mostly lacking. We present GraphCompass, a comprehensive set of omics-adapted graph analysis methods to quantitatively evaluate and compare the spatial arrangement of cells in samples representing diverse biological conditions. GraphCompass builds upon the Squidpy spatial omics toolbox and encompasses various statistical approaches to perform cross-condition analyses at the level of individual cell types, niches, and samples. Additionally, GraphCompass provides custom visualization functions that enable effective communication of results. We demonstrate how GraphCompass can be used to address key biological questions, such as how cellular organization and tissue architecture differ across various disease states and which spatial patterns correlate with a given pathological condition. GraphCompass can be applied to various popular omics techniques, including, but not limited to, spatial proteomics (e.g. MIBI-TOF), spot-based transcriptomics (e.g. 10x Genomics Visium), and single-cell resolved transcriptomics (e.g. Stereo-seq). In this work, we showcase the capabilities of GraphCompass through its application to three different studies that may also serve as benchmark datasets for further method development. With its easy-to-use implementation, extensive documentation, and comprehensive tutorials, GraphCompass is accessible to biologists with varying levels of computational expertise. By facilitating comparative analyses of cell spatial organization, GraphCompass promises to be a valuable asset in advancing our understanding of tissue function in health and disease.

## 1. Introduction

The spatial arrangement and interactions of cells under different physiological and pathological states provide insights into the underlying mechanisms of tissue function and disease progression. Understanding cell spatial organization is not only essential for deciphering physiological processes but also for advancing diagnostic and therapeutic strategies [Rao et al., 2021, Palla et al., 2022a, Williams et al., 2022].

Spatial omics have emerged as a powerful technology for profiling cellular phenotypes in their tissue context. Spatial transcriptomics methods such as 10x Genomics Visium [Ståhl et al., 2016] and Stereo-seq [Chen et al., 2022], as well as spatial proteomics methods such as CODEX [Goltsev et al., 2018] and MIBI-TOF [Keren et al., 2019], can measure molecular profiles while maintaining information about the locations of cells, therefore enabling the study of cell-cell communication [Fischer et al., 2023] and tissue architecture [Fischer et al., 2022, Wu et al., 2022]. Spatial omics technologies have been increasingly leveraged by researchers interested in delineating mechanisms that disrupt tissue homeostasis and cellular niches in diseased individuals. For example, spatial transcriptomics data has been instrumental in deciphering spatial dysregulation in ischemic hearts [Kuppe et al., 2022]. Additionally, spatial proteomics data has been used to elucidate cellular neighborhoods associated with disease progression and response to therapy in breast cancer [Risom et al., 2022].

Related work has looked into identifying cell interactions [Fischer et al., 2023], spatial clusters [Zhao et al., 2021, Varrone et al., 2023], and niche composition in individual samples [Bernstein et al., 2023]. However, methods to compare spatial organization across different sample groups are still lacking. Such methods would be instrumental in elucidating how the arrangement of cell types influences the overall state of a tissue. In this work, we model spatial omics samples as graphs of cells to enable differential analysis of phenotypes. We focus on providing easy-to-use graph metrics and statistical methods for the comparative analysis of cell spatial organization. Studying changes in niche composition and tissue architecture is essential to unlock new insights into the role of tissue organization in prognosis and diagnosis [Rao et al., 2021, Palla et al., 2022a, Williams et al., 2022].

We introduce GraphCompass (**Graph Comp**arison Tools for Differential **A**nalyses in **S**patial **S**ystems), a Python-based framework that brings together a robust suite of graph analysis and visualization methods, specifically tailored for the analysis of cell spatial organization using spatial omics data. Developed on top of `Squidpy` [Palla et al., 2022b] and `AnnData` [Virshup et al., 2021], our methods are easily integrated into existing spatial omics analysis workflows. The framework’s modular design ensures adaptability and compatibility with various single-cell data analysis packages (Figure 1a). Available for community use and collaboration, GraphCompass can be accessed at https://github.com/theislab/graphcompass/, where we provide extensive function documentation and tutorials. We adapted the methods to make them flexible enough to handle large feature spaces (*>*20,000 genes), different resolutions (e.g. spot or single-cell), and multiple modalities of spatial omics data (Figure 1b). To showcase the broad applicability of the methods in our suite, we curate datasets from three different spatial omics techniques and show that our methods recapitulate experimental results, additionally providing novel insights into the global changes in tissue organization under different disease and developmental stages. The collection of omics-adapted methods we present are an effective hypothesis-generating tool that may inform the development of new diagnostic methods and therapeutic targets.

**Fig. 1.**
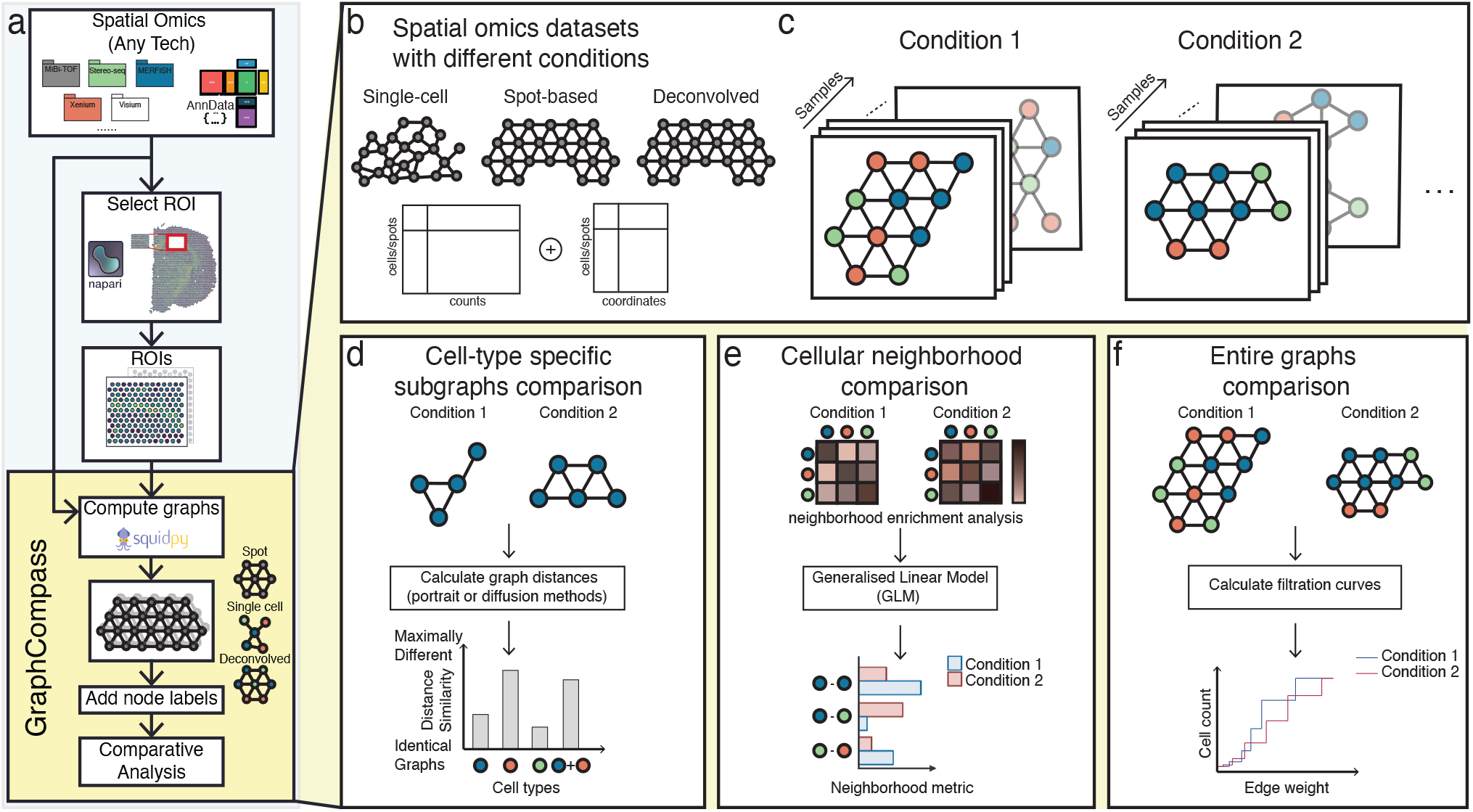
GraphCompass offers graph and statistical analysis methods to compare the spatial organization of cells across different conditions. **a** GraphCompass workflow. All spatial omics datasets that are stored as AnnData objects are currently supported. Support for SpatialData objects [Marconato et al., 2023] will be added in the near future. Select a region of interest (ROI) with napari (GitHub) or use the entire tissue section. We use Squidpy to encode spatial omics measurements as graphs. If available, add node labels, such as cell types. Then, compare graphs across conditions or samples using any of the methods implemented in GraphCompass. **b** The example datasets covered here represent various technologies and different modalities. **c** In our framework, samples are represented as cellular graphs in which nodes correspond to cells or spots and edges denote spatial proximity. Nodes may be labeled (colored) based on their cell type and samples representing the same condition are grouped together to account for sample variation. **d-f** GraphCompass integrates multiple spatial metrics to find statistically significant differences in spatial organization across experimental conditions, utilizing spatial information at various abstraction levels. **d** Analyse graphs that consist of a single cell type and compare them between conditions using graph distance metrics (cell-type-specific subgraphs comparison). **e** Perform neighborhood analysis by retrieving cell-type neighbors enriched in one condition compared to another (cellular neighborhood comparison). **f** Using a holistic approach, compare entire graphs representing data obtained under two or more conditions (entire graphs comparison).

To the best of our knowledge, GraphCompass is the first method to enable differential analysis of spatial organization across conditions at three levels of abstraction: cell-type-specific subgraphs (Figure 1d), multi-cell niches (Figure 1e), and entire graphs (Figure 1f). Though other methods, such as CellCharter [Varrone et al., 2023] and MENDER [Yuan, 2024], also attempt to differentiate samples based on their neighborhood composition, they rely on clustering algorithms, and hence a well-chosen number of clusters. Here, we propose to perform differential niche analysis by studying enriched pairs of neighbor cells. We also present approaches that have never been applied to spatial omics before, such as the Wasserstein Weisfeiler-Lehman kernel and filtration curves. We adapt them to large continuous feature spaces, a typical characteristic of spatial omics data, and show that these metrics are powerful tools to compare samples and sample groups, capturing both local and global information. In this manuscript, we demonstrate the capacity of our methods to reproduce results consistent with previously published findings, as well as provide novel mechanistic hypotheses. To date, GraphCompass is the most comprehensive toolkit aimed at differential neighborhood composition and spatial organization analysis in the context of spatial omics technologies applied to disease studies. We hope that this framework will empower significant advancements in understanding the complexities of cell organization within the spatial context of tissues, both in health and disease.

## 2. Methods

### 2.1 Graph construction

In the realm of spatial omics, nodes in the graph represent either individual cells or predefined spots. The nature of the node depends on the type of spatial omics technology used:

1. Cell-Based Data: For single-cell resolution techniques, each node corresponds to an individual cell. Every node is associated with a node attribute, namely the cell’s transcriptomic profile.
2. Spot-Based Data: Technologies like Visium provide data at the level of spots, which are predefined, regularly spaced areas on a tissue section, each containing multiple cells. In this scenario, each spot, with its aggregated gene expression information, forms a node.

Edges represent spatial proximity between nodes. The edge construction method depends on the data’s layout:

1. Grid Layout: In spot-based technologies like Visium, where spots are arranged in a fixed grid, edge construction is relatively straightforward. Graph edges are typically defined based on direct neighbors in this grid, leading to a structured, regular graph topology.
2. Irregular Layouts: For data not laid out in a grid, defining node adjacency requires more sophisticated methods. Delaunay triangulation is a common approach used here. It involves creating a triangulated mesh such that no node lies inside the circumcircle of any triangle. This method effectively captures the proximity between irregularly spaced cells.

Once the graph is constructed, it serves as a foundational structure for various differential analyses: comparing cell-type-specific subgraphs, cellular neighborhoods, and entire graphs between experimental conditions, developmental stages, or disease states. We use existing methods within Squidpy to compute a spatial graph from various types of spatial omics data. These graphs serve as the input for the analysis and visualization algorithms implemented in our package. We describe these analysis functions in the next sections.

### 2.2. Comparing cell-type-specific subgraphs

We introduce two graph distance metrics to compare cell-type-specific graphs between different conditions: portrait and diffusion methods. Both methods provide similarity scores between two networks of cells that represent two different conditions.

#### 2.2.1. Portrait Method

This method creates a so-called “portrait” of a graph, which is a way to represent the overall structure of the graph [Bagrow and Bollt, 2019]. The portrait of a graph typically includes information about the distribution of distances between nodes and degree distribution. The idea behind the portrait method is to capture the essence of the graph’s topology in a comprehensive snapshot.

Given two graphs *G* and *G*^*′*^, we first define the network portrait *B* of each graph as an array with *l* × *k* elements, such that

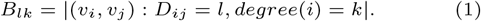

Here, (*v*_*i*_, *v*_*j*_) are node pairs of graph *G* such that the shortest path between *v*_*i*_ and *v*_*j*_, *D*_*ij*_, equals *l*. The degree of a node is defined as the number of edges incident to that node. We do not compare *G* and *G*^*′*^ directly. Instead, we compare their network portraits *B* and *B*^*′*^, such that △(*G, G*^*′*^) ≡ △(*B, B*^*′*^) (that is, such that the difference between the network portraits approximates the difference between the graphs). To compare the network portraits, we calculate the weighted distributions *P* (*k, l*) and *Q*(*k, l*), such that

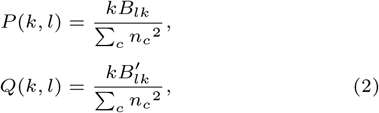

where *nc* represents the number of nodes within a given connected component *c*, and Σ_*c*_ *n*_*c*_ = *N*, with *N* being the total number of nodes in the graph. We subsequently compare the two distributions using the Jensen-Shannon divergence:

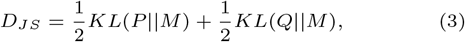

where *KL* is the Kullback–Leibler divergence and 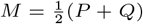, *D*_*JS*_ is the similarity score between the cell-type-specific graphs *G* and *G*^*′*^, each representing a different experimental condition or co-variate. Here, a high similarity score implies maximally different graphs, and a low score implies that graphs are nearly identical. This comparison is repeated for every cell type present in both graphs. Cell-type-specific similarity scores are jointly visualized to determine which cell types are most similarly organized across both conditions.

#### 2.2.2. Diffusion Method

Diffusion, in the context of graphs, refers to the process of spreading a certain amount of an imaginary substance (like information, heat, etc.) across the nodes of a graph over time. Diffusion on graphs can be intuitively understood through the analogy of balls connected by springs. When you impart energy to one ball in the system (for example, by hitting or pushing it), this energy is represented by the ball’s movement. As the ball starts moving, it stretches or compresses the springs connected to it (the edges in the graph). This, in turn, transfers energy to the balls (nodes) at the other ends of these springs. Balls directly connected to the moving ball receive the energy first, and then the energy propagates to others in a ripple-like effect. The overall structure of the graph (how balls are connected by springs) affects the energy diffusion pathway and rate.

Diffusion on graphs is implemented by NetLSD [Tsitsulin et al., 2018], a Python library that encodes a so-called “trace signature” to capture the energy diffusion process. The trace signature is computed as follows: Given graph *G*, calculate its normalized Laplacian as

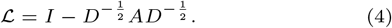

*A* and *D* are the adjacency and degree matrix of *G*, respectively. Next, we compute the closed-form solution to the heat equation associated with the normalized Laplacian, which is defined as

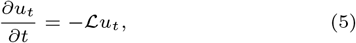

where *u*_*t*_ represents the imaginary “energy” of a given node at time *t*. The solution to the heat equation is then computed as

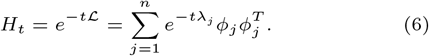

Here, *H*_*ij,t*_ quantifies the amount of energy transferred from node *v*_*i*_ to node *v*_*j*_ at time *t*. As a last step, we compute the heat trace *h*_*t*_ as the trace of *H*_*t*_, such that

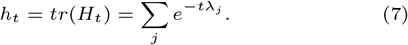

To compare two graphs, we simply compute the *L*_2_ distance between the corresponding heat traces computed at different times *t*,

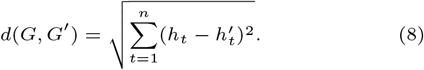

### 2.3. Comparing cellular neighborhoods

To compare cellular neighborhoods between different conditions, we leverage Generalized Linear Models (GLMs). These models allow us to determine statistically significant changes in the neighborhood enrichment (i.e., the enrichment of spatial proximity between two cell types) across multiple conditions, offering a deeper understanding of the spatial density and distribution of specific cell types relative to others under a given condition. Here, we refer to a pair of cell types as “enriched” if they neighbor each other more often than we would expect based on random chance. We first compute neighborhood enrichment in each sample separately using `Squidpy’s nhood enrichment` function. This function calculates the observed number of each cell type pair, which is then compared against the expected frequency. This expected frequency is determined through permutation tests.

The `nhood enrichment` function returns a *n* × *n* matrix *Z* containing enrichment z-scores. *Z*_*ij*_ represents the enrichment of the pair that consists of cell type *i* and cell type *j*. Since this matrix is symmetric, we extract the upper triangular portion, which we flatten to obtain a 1 × *n* vector representing neighborhood enrichment in a single sample. Given *m* samples, we concatenate their corresponding vectors along the condition axis to obtain a *m* × *n* matrix for usage in linear models (neighborhood enrichment z-scores) or Poisson and Quasi-Poisson Generalised Linear Models. For the latter two, we model the counts of each observed cell type pair in each neighborhood rather than the z-scores. The Quasi-Poisson model is particularly appropriate when neighborhood counts are sparse for a pair of cell types. We include a fixed linear term to account for the “batch/subject/patient” co-variate, and an interaction term between all levels of the condition and cell type pair factors to fit the models.

### 2.4. Comparing entire graphs

We present two methods to perform holistic graph comparisons: filtration curves and Weisfeiler-Lehman graph kernels. Both methods result in graph embeddings that can be compared against one another to obtain a broad measure of tissue architecture similarity.

#### 2.4.1. Filtration Curves

In the context of Topological Data Analysis (TDA), filtrations are a fundamental concept used to understand the shape of data [O’Bray et al., 2021]. The basic idea is to gradually “grow” or “filter” the data and observe how topological features such as connected components, holes, and voids evolve. We define a graph filtration as a sequence of nested subgraphs ∅ ⊆ *G*_1_ ⊆ *G*_2_ … *G*_m_ ⊆ *G*, ordered by edge weights. Let *G* = (*V, E, w*) be a weighted graph, where *w* : *E* → ℝ is the weight function assigning a real number to each edge, here defined as the Euclidean distance between the gene expression matrices associated with neighboring nodes. To generate the filtration curve, we order the edges based on their weights, obtaining a series of weights *w*_1_ ≤ *w*_2_ … *w*_*m*−1_ ≤ *w*_*m*_. O’Bray et al. [2021] define the *i*^th^ graph in the filtration, *G*_*i*_, as the subgraph that includes all edges whose weight is less than or equal to *w*_*i*_ as well as all nodes connected by said edges. Since our distance function can take on any positive real number, we compute ten threshold values from the collection of edge weights to restrict the algorithm’s computation time. We define the threshold values as the 10th, 20th, …, 90th and 100th percentile. At every filtration step, the algorithm analyzes the properties of the subgraph by evaluating a graph descriptor function. Assuming every node has been assigned a node label (i.e., a cell type), we can simply compute the number of each cell type present in the subgraphs. Computing and comparing filtration curves is an efficient approach for representing graphs and contrasting two graphs or sets of graphs.

#### 2.4.2. Weisfeiler-Lehman Graph Kernels

The Weisfeiler-Lehman (WL) graph kernel is a powerful technique used in graph theory and machine learning, particularly in the context of graph classification and similarity analysis. Boris Weisfeiler and Andrei Lehman introduced it in the late 20th century as a graph isomorphism test [Weisfeiler and Leman, 1968]. Though it has been shown that there are non-isomorphic graphs that cannot be distinguished by this algorithm, it has been successfully implemented as a graph similarity measure [Shervashidze et al., 2011]. Broadly speaking, the algorithm consists of three steps: node label augmentation, iteration, and kernel computation. In each iteration, the node label of a given node is transformed into an augmented label, or multi-set of labels, that contains the original label as well as the labels of the given node’s neighbors. The augmented label is subsequently hashed, resulting in a new, compressed node label. Given a graph *G* = (*V, E*), where *V* is a set of nodes (vertices) and *E* is a set of edges, we can define the node label augmentation step as

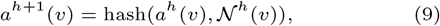

We define *a*^*h*^(*v*) as the compressed label of node *v* at iteration *h*. Similarly, 𝒩^*h*^(*v*) represents the neighbor labels at iteration *h*. Lastly, we define *a*^0^(*v*) as the original label of node *v*. The node labeling step is repeated for a pre-specified number of iterations. After the iteration process, the labels assigned to the nodes are used to compute a kernel matrix. This matrix quantifies the structural similarity between pairs of graphs. The original formulation of the algorithm restricts its use to graphs with discrete labels. However, some of the more common spatial omics methods, most notably Visium, do not produce single-cell-resolved data. Each spot may contain more than one cell, complicating cell type assignment. The spot is best represented by its associated gene expression matrix. The Wasserstein WL kernel [Togninalli et al., 2019] extends the WL kernel from the discrete to the continuous case. We define *a*^*h*^(*v*) as the attribute of node *v* at iteration *h*. Let *w*(*v, u*) be the weight of the edge between nodes *v* and *u*. Then, the updated node attribute at iteration *h* + 1 is computed as

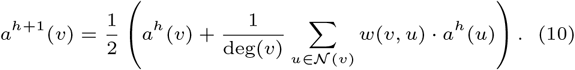

Once node embeddings have been computed, the algorithm evaluates the distance between pairs of nodes 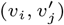 for each *v*_*i*_ ∈ *V* and each 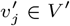, resulting in distance matrix 𝒟. Here, we define the distance between nodes *v*_*i*_ and 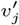 as the Euclideandistance between their corresponding gene expression matrices. Lastly, the algorithm quantifies the similarity of graphs *G* and *G*^*′*^ by measuring the Wasserstein distance between them as

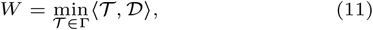

where 𝒯 ∈ Γ is a transport matrix and ⟨·, ·⟩ is the Frobenius dot product.

## 3. Results

In the next sections, we demonstrate the utility of GraphCompass methods by analyzing three datasets derived from three different technologies and spatial systems (MIBI-TOF breast cancer [Risom et al., 2022], 10x Genomics Visium heart [Kuppe et al., 2022], and Stereo-seq axolotl [Wei et al., 2022]). We only use analysis and visualization functions implemented in GraphCompass, highlighting what can be learned from each function.

### 3.1. Spatial organization of the tumor microenvironment and breast cancer progression

Risom et al. [2022] used multiplexed ion beam imaging by time of flight (MIBI-TOF) [Keren et al., 2019] with a 37-plex antibody staining panel to study changes in the tumor microenvironment during the transition from ductal carcinoma in situ (DCIS) to invasive breast cancer (IBC), allowing them to identify spatial and functional changes in various cell types, including myoepithelial cells, fibroblasts and immune cells (Figure 2a). They compared normal samples to both DCIS samples and IBC patient samples. DCIS samples can be further divided into progressors (samples that progress from DCIS to IBC) and non-progressors. The subset of data we analyze consists of 67 samples (*N*_*Normal*_ = 9, *N*_*Progressors*_ = 14, *N*_*Non*−*progressors*_ = 44). As part of their effort to identify features that distinguish transitioning samples from non-transitioning samples, the authors used a masking approach to gauge the thickness and continuity of the myoepithelial barrier in multiplexed images. An important, yet surprising, finding of this experiment is that myoepithelial disruption occurs in lesions that did *not* become invasive (non-progressors), while the myoepithelium of DCIS patients that *do* develop IBC (progressors) stayed mostly intact. A robust myoepithelial barrier is a key feature of healthy breast tissue, meaning that progressor samples more closely resemble normal breast samples in terms of myoepithelial robustness than non-progressor samples do. Risom et al. [2022] suggest that myoepithelial disruption may be a protective mechanism against progression to invasive cancer.

**Fig. 2.**
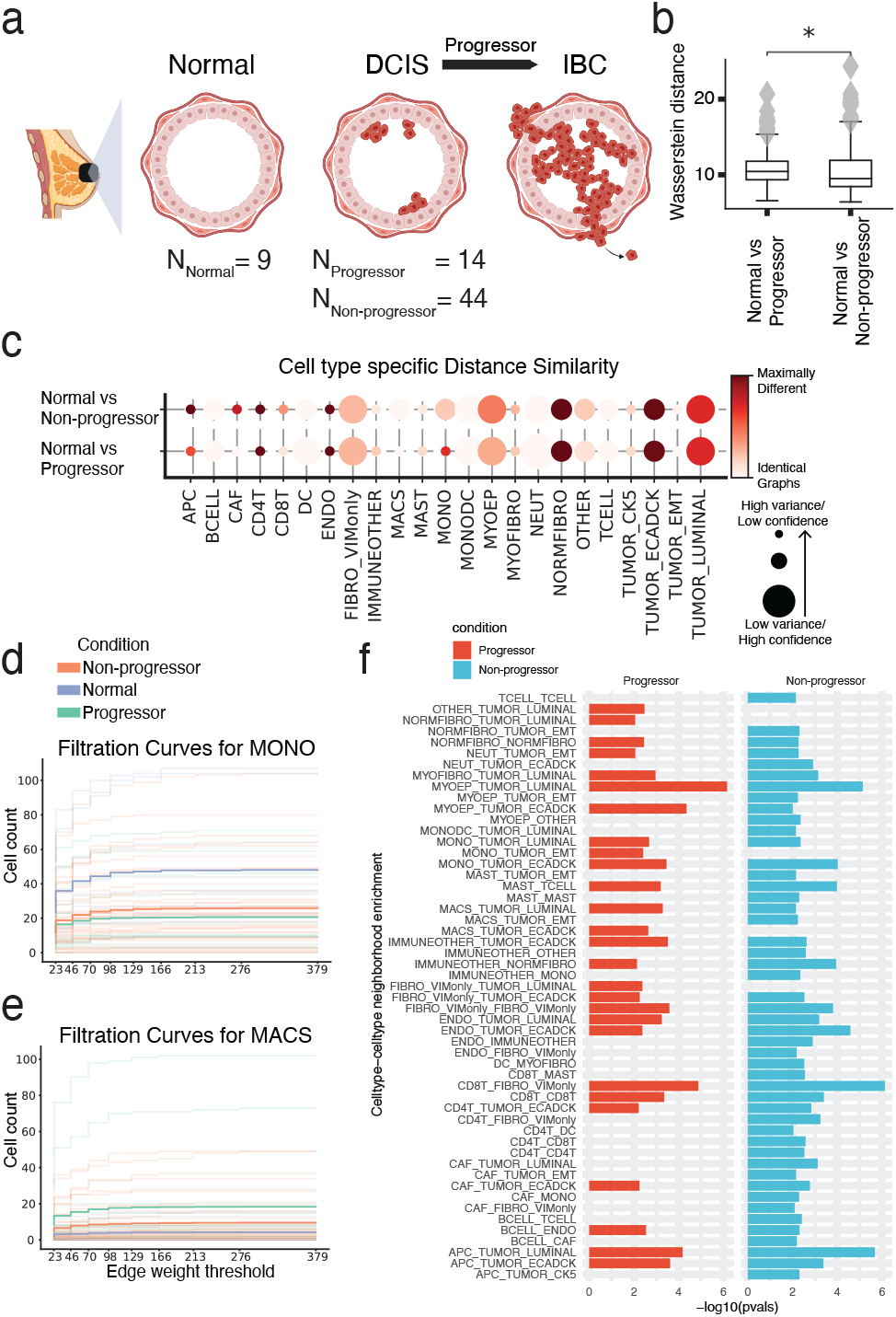
MIBI-TOF dataset studying the role of the tumor microenvironment (TME) in breast cancer progression. **a** Schematic figure describing the different biological conditions investigated in this study. **b** Comparing entire tissue samples, using Weisfeiler-Lehman Graph Kernels, to show the overall similarity in spatial organization across two conditions (normal versus non-progressors and normal versus progressors). The smaller the Wasserstein distance, the more similar the spatial organization is under the two compared conditions. **c** Cell-type-specific subgraphs comparison, using the portrait method, across condition pairs (normal versus non-progressors and normal versus progressors). The size of the dot is indicative of the similarity score variance over samples. The larger the dot size, the lower the score variance and the higher the score confidence is. **d, e** Filtration curves (Normal, Non-progressor and Progressor) for **d** Monocytes and **e** Macrophages. We plot a filtration curve for every sample, as well as the mean curve for every condition, which can be identified by the thicker, darker lines. Large vertical steps towards the left of the plot indicate low density, whereas large vertical steps towards the right of the plot indicate high density. **f** Enrichment of various cell type pairs in progressors and non-progressors, where the control condition acts as a baseline.

We employed GraphCompass to further investigate the downstream effects of myoepithelial disruption on breast tissue architecture at different scales. We first used a holistic approach, Weisfeiler-Lehman Graph Kernels (Section 2.4.2), to assess the overall similarity between the architecture of normal breast tissue and the spatial organization of non-progressor and progressor samples. Based on this holistic view of breast cellular organizational structure, we find that normal tissue resembles non-progressor samples more closely than progressor samples (Figure 2b). Next, we generated cell-type-specific subgraphs using the portrait distance method (Section 2.2.1). These subgraphs indeed suggest that the spatial organization of myoepithelial cells (MYOEP) in normal breast tissue is significantly more similar to that in progressor tissue than that in non-progressor tissue (*p* = 7.9*e*^−4^, Student’s t-test comparing similarity score means between I. normal vs progressor and II. normal vs non-progressor) (Figure 2c). GraphCompass was thus able to confirm the previously reported finding that non-progressor tissue is characterized by its compromised myoepithelial layer, distinguishing it from healthy and progressor tissue.

To further attempt to explain the protective quality of the disintegrating myoepithelial barrier, we executed a neighborhood analysis (Section 2.3) to determine which types of cells are more likely to co-occur in non-progressor samples than in progressor samples and normal breast samples. Interestingly, immune cells were more likely to neighbor other immune cells in non-progressor samples compared to normal breast samples, indicating that non-progressors mount an immune response to the tumor, recruiting T lymphocytes (TCELL), B lymphocytes (BCELL), and dendritic cells (DC) to the site of the tumor. Indeed, CD4T-CD4T, CD4T-CD8T, B cell-T cell, and CD4T-DC were all enriched in non-progressor samples compared to normal samples (Figure 2f). Notably, we did not observe an enrichment of these neighbor pairs in progressor samples. We hence hypothesize that a thinner myoepithelial barrier protects against the transition to invasive breast cancer by contributing to the development of a “hot” tumor, i.e., a tumor that presents with a microenvironment characterized by heightened immune activity, often featuring tumor-infiltrating lymphocytes [Duan et al., 2020]. The “temperature” of immune environments has indeed been shown to play a crucial role in shaping the trajectory of disease progression from pre-invasive lesions to invasive cancer [Galon et al., 2010, Fridman et al., 2017]. The compromised myoepithelial barrier in non-progressor samples may allow immune cells, particularly T lymphocytes, greater access to the tumor microenvironment, increasing their presence around tumors. Our analysis suggests that these tumor-infiltrating T cells may eventually trigger cancer cell death, preventing progression to invasive breast cancer.

In addition, we found that monocyte (MONO) organization in normal tissue is more similar to monocyte organization in non-progressors than that in progressors (*p* = 7.6*e*^−3^, Student’s t-test comparing similarity score means between I. normal vs. progressor and II. normal vs. non-progressor) (Figure 2c). Furthermore, the filtration curves (Section 2.4.1) show that the average number of macrophages (MACS) is higher in progressor samples than in non-progressors and control samples (Figure 2e). In mouse models of cancer, monocytes have been observed to migrate to the site of the tumor, where they differentiate into tumor-associated macrophages (TAMs). Multiple independent breast cancer studies have identified the TAM signature and density as markers of tumor progression [Lin et al., 2003, Arwert et al., 2018, Cassetta et al., 2019]. Our results suggest that progressor monocytes have differentiated into macrophages, which may affect progressor prognosis. We could not establish differences in the organization of luminal tumor cells between progressors and non-progressors (Figure 2c). Therefore, tumor spatial organization neither seems to cause nor appears to be immediately affected by myoepithelial integrity.

Understanding and manipulating the immune environment is essential for developing targeted therapeutic strategies to enhance immune responses and restrain cancer progression. Though further experimental validation is beyond the scope of our manuscript, we have shown that the algorithms implemented in GraphCompass generate results consistent with previously published findings, namely that myoepithelial barrier disintegration is associated with favorable disease outcomes. We have also demonstrated the use of GraphCompass as a hypothesis-generating tool, offering a potential explanation as to why myoepithelial loss protects against tumor progression.

### 3.2. Myocardial tissue reorganization following ischemic injury

Kuppe et al. [2022] conducted a comprehensive study to examine the changes that occur in the cardiac transcriptome and epigenome following a heart attack. They integrated data from three different modalities: single-cell RNA-seq, chromatin accessibility data, and spatial transcriptomics data generated by the Visium platform [Ståhl et al., 2016]. Their data contains samples from patients who experienced myocardial infarction and healthy individuals. Samples were taken from different physiological zones of the myocardium (RZ, remote zone; BZ, border zone; IZ, ischaemic zone; FZ, fibrotic zone) (Figure 3a). Here, we focus on the experiments that were based on spatial transcriptomics data. These experiments show changes in the organization of cardiomyocytes and myeloid cells after ischemic injury.

**Fig. 3.**
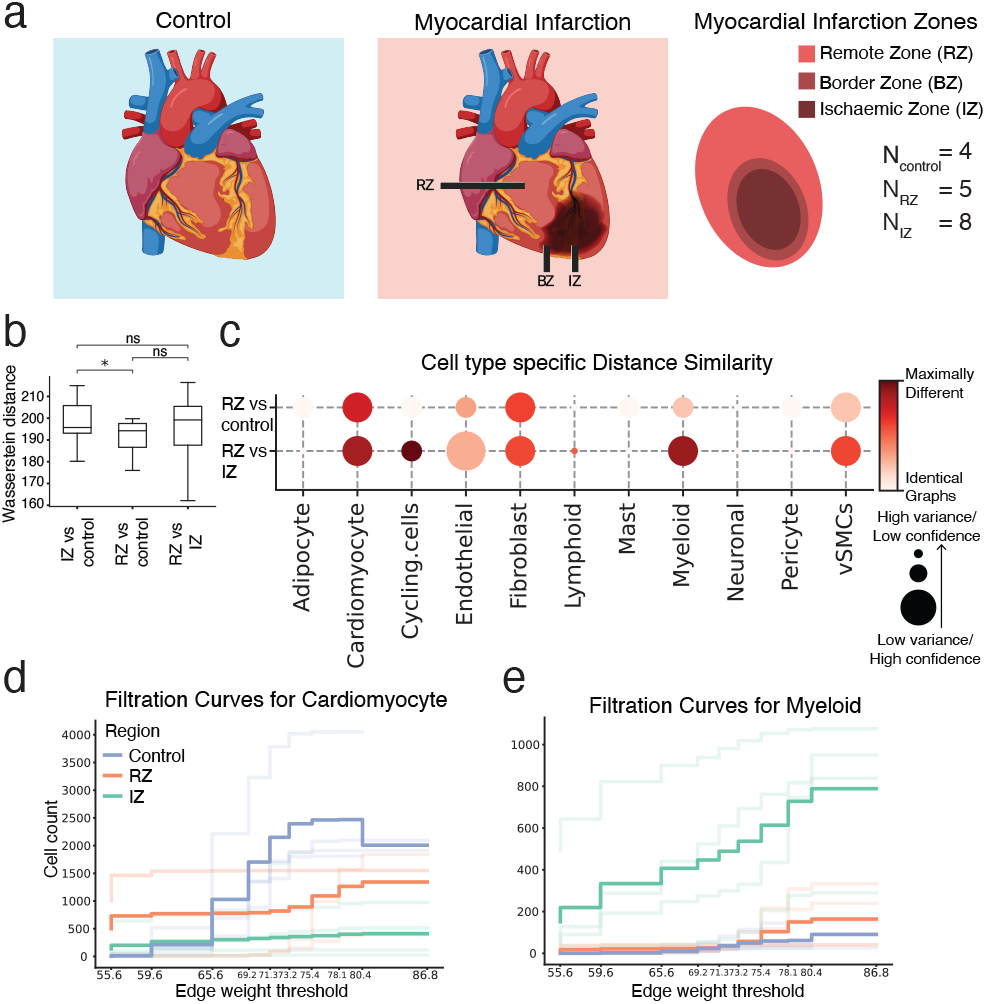
Visium dataset studying myocardial tissue architecture following ischemic injury. **a** Schematic figure describing the different physiological zones studied: the ischaemic zone (IZ), border zone (BZ), the unaffected left ventricular myocardium (remote zone, RZ), and control samples. **b** Comparing entire tissue samples, using Weisfeiler-Lehman Graph Kernels, to show the overall similarity in spatial organization across two conditions (RZ versus control and RZ versus IZ). The smaller the Wasserstein distance, the more similar the spatial organization is under the two compared conditions. **c** Cell-type-specific subgraphs comparison, using the portrait method, across condition pairs (RZ versus control and RZ versus IZ). The size of the dot is indicative of the similarity score variance. The larger the dot size, the lower the score variance and the higher the score confidence is. **d, e** Filtration curves (RZ, IZ and Control) for **d** Cardiomyocytes and **e** Myeloid cells. We plot a filtration curve for every sample, as well as the mean curve for every condition, which can be identified by the thicker, darker lines. Large vertical steps towards the left of the plot indicate low density, whereas large vertical steps towards the right of the plot indicate high density.

To study the effect of ischemic injury beyond the initial site of the injury, we performed a comparison of samples taken from three physiological regions: the IZ, the unaffected left ventricular myocardium (RZ), and control cardiac tissue (*N*_*IZ*_ = 8, *N*_*RZ*_ = 5, *N*_*Control*_ = 4). We focused our analysis on these three regions to better understand whether RZ is affected by the ischemic injury and therefore more similar to the IZ or is protected from the injury and thus more similar to control tissue. Using the entire graph comparison approach (Section 2.4.2), we show that the spatial arrangement of the RZ is not significantly more similar to the arrangement of the IZ than that of the control (Figure 3b). This indicates that the remote zone might not be impacted, or only partially impacted, by the myocardial infarction. To further study the effects of ischemic injury at the cellular organization level, we utilized the cell-type-specific portrait method (Section 2.2.1). We found that the organization of cardiomyocytes in the RZ differed from that in the normal tissue samples. It also differed from cardiomyocyte organization in the IZ. Overall, the spatial arrangement of cardiomyocytes in the RZ is slightly more similar to the arrangement in the control samples than to the arrangement in the IZ, though the effect is not significant (*p* = 0.25, Student’s t-test comparing similarity score means between I. RZ vs control and II. RZ vs IZ) (Figure 3c). This finding indicates that the cardiomyocytes in the remote ventricular myocardium are impacted by the injury, though to a lesser extent than the cardiomyocytes in the IZ. Our results also suggest that the arrangement of myeloid cells in the RZ is significantly more similar to that in the control tissue than that in the IZ (*p* = 2.2*e*^−7^, Student’s t-test comparing similarity score means between I. RZ vs control and II. RZ vs IZ) (Figure 3c). This supports the notion that the damage inflicted by ischemic injury on myeloid cells is localized at the injury site. The filtration curves also show that cardiomyocyte organization in the RZ is affected by the injury (Figure 3d), while myeloid organization is not (Figure 3e). In particular, the curves show that both the number and density of cardiomyocytes in the RZ have been impacted by the infarction.

Collectively, our results support the finding that myocardial infarction can have localized or systemic impacts on different cell types. Though damage typically originates in a specific area of the heart, we observe that the consequences can extend beyond the initial site of injury. Indeed, experimental studies have suggested that the size of the infarct depends on the post-infarct inflammatory response [Frangogiannis, 2014].

### 3.3. Restoration of axolotl brain function upon injury: Comparing healthy and regenerated brains

Wei et al. [2022] used the Stereo-seq technology [Chen et al., 2022] to generate spatial omics data spanning six axolotl developmental stages and seven regeneration phases. The axolotl is a type of salamander, known for its remarkable ability to regenerate lost body parts. This ability makes them an invaluable model for studying tissue regeneration and wound healing, potentially offering insights applicable to human medicine (Figure 4a). To shed light on the molecular events that precede regeneration, the authors removed a part of the brain and then collected spatial transcriptomics data 2, 5, 10, 15, 20, 30, and 60 days post-injury. They claim that 60 days post-injury, brain cell composition and the spatial distribution of cell types are restored.

**Fig. 4.**
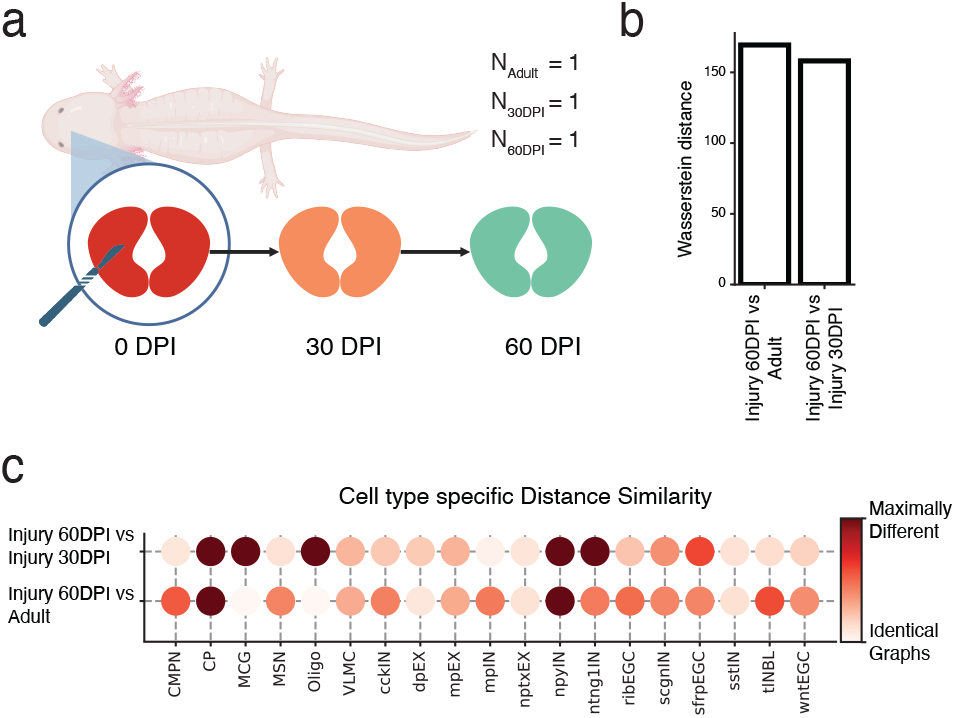
Stereo-seq dataset studying the axolotl brain during development and regeneration. **a** Schematic figure describing the subset of regeneration stages we investigated. On the day of injury (0 days post-injury, DPI), a section of the brain was removed. We compared a tissue sample collected 30 days post-injury (30 DPI) with a section obtained 60 days post-injury (60 DPI) and a control sample from an unharmed adult axolotl. **b** Comparing entire tissue samples, using Weisfeiler-Lehman Graph Kernels, to show the overall similarity in spatial organization across two stages (Injury 60 DPI versus Adult and Injury 60 DPI versus Injury 30 DPI). The smaller the Wasserstein distance, the more similar the spatial organization is under the two compared conditions. **c** Cell-type-specific subgraphs comparison, using the portrait method, across condition pairs (Injury 60 DPI versus Adult, Injury 60 DPI versus Injury 30 DPI).

To assess tissue restoration success, we focused on studying the last two regenerative stages using two samples collected 30 and 60 days post-injury (30 DPI and 60 DPI). We compared the 60 DPI sample against the 30 DPI sample as well as a control adult sample from the development data set (*N*_30*DP I*_ = 1, *N*_60*DP I*_ = 1, *N*_*Adult*_ = 1). The aim of our analysis is to understand whether, after 60 days, the regenerating axolotl brain is more similar to the unharmed adult brain or the 30 DPI brain. If the axolotl brain has indeed completely regenerated, we would expect to see that both the distribution of cell types and their spatial organization have been restored, mimicking that of the control adult sample. Comparing the 30 DPI, 60 DPI, and control sections at the sample level (Section 2.4.2), we show that the 60 DPI brain is slightly more similar to the 30 DPI brain than to the adult brain, indicating that the arrangement of cells has not been fully restored post-injury (Figure 4b), though the differences are subtle.

Comparison of the cell-type-specific subgraphs further supports our conclusion that the spatial organization of the regenerated brain differs from the organization of the healthy brain. Indeed, the portrait graph (Section 2.2.1) indicates that the organization of multiple cell types in the 60 DPI sample resembles the 30 DPI organization more so than the adult brain organization. For example, one cell type that is arranged similarly in the 30 DPI and 60 DPI samples is the telencephalon neuroblast (tlNBL), which has been shown to have a role in telencephalon neurogenesis during regeneration [Lust et al., 2022] (Figure 4c), indicating that regeneration may not yet be complete 60 days after the injury. However, the portrait plot also shows several cell types in the 60 DPI sample whose spatial organization is similar to that of adult cells. These cell types include dorsal pallium excitatory neurons (dpEX) and Sfrp+ ependymal glial cells (sfrpEGC). This suggests that the arrangement of dpEX and sfrpEGC cells is restored 60 days post-injury.

Wei et al. [2022] observe that development and regeneration are characterized by many of the same processes, including neuronal differentiation and migration, but that several pathways were uniquely upregulated in regenerating brains. In addition, they identify two subtypes of ependymoglial cells (EGCs), one of which is present in the developing brain, while the other is found only in the regenerating brain. It is possible that these biological differences underlie the incomplete restoration of cell spatial organization in regenerating brains, but more data is needed to draw robust conclusions.

To summarize, we find that the arrangement of some cell types is successfully restored in the 60 days following brain injury. However, we also highlighted differences in the organization of the 60 DPI brain and the healthy adult brain, indicating that the former had not been fully regenerated at the 60-day mark.

## 4. Discussion

GraphCompass is a comprehensive graph analysis framework that provides quantitative methods to compare cell spatial organization across physiological systems, pathological states, and developmental stages. Compatible with spatially resolved transcriptomics and proteomics data, GraphCompass integrates multiple graph-based and statistical approaches for investigating spatial graphs at three different levels of abstraction: individual cell types, multi-cell neighborhoods, and entire samples. These methods were adapted for spatial omics data, such that they can handle high-dimensional features, flexible node identities (spot or single cell) and variable edge weight definitions.

Differences in cell spatial organization can be indicators of disease states, or correlate with how patients respond to treatments. Studying cell spatial organization across individuals can provide insights into developmental and regenerative processes, which can guide the development of engineered tissues and organoids. We believe that GraphCompass will significantly advance our understanding of the role of tissue architecture in healthy development, disease onset, and recovery.

In this manuscript, we have demonstrated the capabilities of GraphCompass through its application to datasets derived from diverse technologies, highlighting the biological insights that can be obtained from the various metrics it implements. Developed in Python, GraphCompass interfaces seamlessly with Squidpy and AnnData, enhancing its scalability and the potential for expansion with new methodologies. With GraphCompass, our aim is to offer the computational biology community user-friendly and accessible graph comparison methods, empowering both experimental and computational scientists in the analysis and interpretation of tissue architecture differences across different biological phenotypes.

## 6. Code availability

All spatial metrics described here are implemented in a Python package available at https://github.com/theislab/graphcompass/. Code documentation and tutorial notebooks can also be found on GitHub.

## 7. Data availability

All the data sets used in this work are publicly available through open-access repositories. The Stereo-seq axolotl data [Wei et al., 2022] are available in the Spatial Transcript Omics DataBase (STOmics DB) under https://db.cngb.org/stomics/artista/. The MIBI-TOF breast cancer data [Risom et al., 2022] are available in a public Mendeley data repository: https://data.mendeley.com/datasets/d87vg86zd8. The Visium heart data [Kuppe et al., 2022] are available in the Zenodo data repository under https://zenodo.org/record/6578047.

## Acknowledgments

The authors thank Bastian Rieck for his valuable suggestions. F.J.T. acknowledges support from the European Union (ERC, DeepCell – 101054957). K.H. acknowledges financial support from the Joachim Herz Stiftung via Add-on Fellowships for Interdisciplinary Life Science and support from the Helmholtz Association under the joint research school ‘Munich School for Data Science’.

